# The Calcium Transient Coupled to the L-Type Calcium Current Attenuates Action Potential Alternans

**DOI:** 10.1101/2023.04.25.538350

**Authors:** Mark Warren, Steven Poelzing

**Affiliations:** Fralin Biomedical Research Institute at Virginia Tech Carilion, Center for Vascular and Heart Research. Roanoke, VA; Dept. of Biomedical Engineering and Mechanics at Virginia Tech, Blacksburg, VA; Dept. of Internal Medicine at Virginia Tech Carilion, Roanoke, VA

**Keywords:** Alternans, L-type calcium channel, Action potential, Calcium transient

## Abstract

**Background:** Action potential (AP) alternans are linked to increased arrhythmogenesis. It is suggested that calcium (Ca^2+^) transient (CaT) alternans cause AP alternans through bi-directional coupling feedback mechanisms because CaT alternans can precede AP alternans and develop in AP alternans free conditions. However, the CaT is an emergent response to intracellular Ca^2+^ handling, and the mechanisms linking AP and CaT alternans are still a topic of investigation. This study investigated the development of AP alternans in the absence of CaT.

**Methods:** AP (patch clamp) and intracellular Ca^2+^ (Fluo-4 epifluorescence) were recorded simultaneously from isolated rabbit ventricle myocytes perfused with the intracellular Ca^2+^ buffer BAPTA (10-20 mM) to abolish CaT and/or the L-type Ca2+ channel activator Bay K 8644 (25 nM).

**Results:** After a rate change, alternans were critically damped and stable, overdamped and ceased over seconds, underdamped with longer scale harmonics, or unstably underdamped progressing to 2:1 capture. Alternans in control cells were predominantly critically damped, but after CaT ablation with 10 or 20 mM BAPTA, exhibited respectively increased overdamping or increased underdamping. Alternans were easier to induce in CaT free cells as evidenced by a higher alternans threshold (ALT-TH: at least 7 pairs of alternating beats) relative to control cells. Alternans in Bay K 8644 treated cells were often underdamped, but the ALT-TH was similar to control. In CaT ablated cells, Bay K 8644 prolonged AP duration (APD) leading predominantly to unstably underdamped alternans.

**Conclusions:** AP alternans occur more readily in the absence of CaT suggesting that the CaT dampens the development of AP alternans. The data further demonstrate that agonizing the L-type calcium current without the negative feedback of the CaT leads to unstable alternans. This negative feedback mechanism may be important for understanding treatments aimed at reducing CaT or its dynamic response to prevent arrhythmias.

## INTRODUCTION

Cardiac electrical alternans are an abnormal cardiac impulse oscillatory mode characterized by a period doubling of the action potential’s (AP) stimulus-response pattern such that every-other beat is either a long or a short AP, herein referred to as 2:2 capture. Electrical alternans are clinically relevant because they underlie the development of T-wave alternans (TWA),^1^ an electrocardiographic phenomenon associated to an increased risk of life threatening arrhythmia.^2^ Further, the development of TWA and their progression towards larger magnitude oscillations and/or T-wave multupling (increased complexity T-wave oscillations) preceded ventricular tachyarrhythmias in dogs^3^ and humans.^4^ In patients with atrial flutter, overdrive pacing lead to atrial fibrillation in those hearts developing AP alternans more frequently and during pacing at longer cycle lengths.^5^ Additionally, AP alternans have been implicated as a cellular mechanism driving spiral wave break-up in simulations^6^ and as a mechanism responsible for complex AP dynamics sustaining ventricular fibrillation.^7, 8^

Functionally linked to the development of AP alternans, hearts may also develop cytosolic Ca^2+^ transient (CaT) alternans,^9^ and these in turn mediate the development of mechanical and pulsus alternans.^10–12^ The relationship between TWA, AP alternans, pulsus alternans, and CaT alternans is complex. On the one hand, local analysis of both AP^13–15^ and CaT^14–16^ signals demonstrated that either may develop alternans exhibiting a site-specific phase such that different locations of the myocardium have discordant signals in CaT or AP,^14^ making the translation of the local fluctuations to the organ wide manifestation of the phenomena as fluctuations in T-wave and pulse pressure hard to predict. On the other hand, AP and CaT are functionally bidirectionally coupled,^17, 18^ rendering it hard to determine how alternans in any of the signals influences the development of alternans in its counterpart, and ultimately determining what is the root cause of arrhythmogenic AP alternans.

The development of AP alternans can be mechanistically considered solely from the perspective of the ion currents configuring the AP, involving a cyclical incomplete recovery of one or more of the ion channels configuring the AP.^19^ In this action potential centered scheme, an abrupt shortening of the diastolic interval (DI; for example via an abrupt shortening of the constant pacing cycle length) will lead to an incomplete ion current recovery and a resulting shorter AP. However, the consequence of the shorter AP will be a subsequent longer DI which will then allow for some recovery of time-dependent ion channel gating in this activation cycle such that the next AP in line will be ‘restored’/longer. However, if pacing cycle length has been kept constant through this sequence, the subsequent DI will now again be shorter, setting thus the conditions for a recursive paradigm with an abnormally short AP every-other beat interspersed with a restored AP, i.e. the characteristic long-short-long-short pattern of alternans. In theory the pattern will perpetuate if the alternating extents of ion channel recovery (i.e. alternating incomplete vs complete recovery) can lock into a stable mode of oscillation. It was proposed that such stability is determined by the steepness of AP duration (APD) restitution curve.^20^

In any case, the above described action potential centered scheme does not consider the interaction of the dynamically changing AP signals with the counterpart dynamic changes in CaT, which should, via a well-established bidirectional coupling between the signals, modulate AP alternans and could even theoretically drive the abnormal impulse oscillations. To this point, it was shown that CaT can exhibit alternans in the absence of and preceding AP alternans,^21^ which has driven (to a large extent) the predominant view that CaT alternans are a primary source of cardiac alternans driving AP alternans through a feedback mechanism.^9, 17, 18, 22^

To date, the counterpart scenario in which AP alternans develop in the absence of CaT alternans has not been studied in detail. Verification or invalidation of this scenario is a principle task since it will either reject or support the requirement of CaT alternans for the formation of AP alternans. To this end, previous studies investigated whether AP alternans were altered after pharmacological interventions designed to abolish the CaT and/or CaT alternans were implemented, including the use of caffeine,^11, 23, 24^ ryanodine,^11, 23, 25^ thapsigargin,^26^ ryanodine in combination with thapsigargin,^27^ BAPTA-AM,^27, 28^ or BAPTA.^27^ In all cases, the reported data suggested that AP alternans were either abolished, reduced in magnitude, and/or they required faster activation rates to be elicited.^11, 23–25, 27, 28^

Importantly, the CaT is an emergent phenomenon dependent on numerous intracellular calcium handling mechanisms.^29^ A number of calcium handling mechanisms occur on the membrane, and therefore there is still some question on whether the entire CaT is necessary to drive, or is truly responsible for AP alternans.^29^ Furthermore, AP alternans are temporally complex, and due to experimental limitations are often quantified over short recording intervals. It is therefore unknown whether AP alternans are always driven by CaT, and how sarcolemmal calcium handling proteins agonize or perhaps even attenuate AP alternans. The purposes of this study are to describe in isolated ventricular myocytes the types of AP alternans that arise after a rate change, to demonstrate whether AP alternans can occur in the absence of CaT alternans, to quantify AP alternan responses to agonizing the L-Type calcium channel (I_CaL_), and to determine the role of I_CaL_ in the regulation of AP alternans. We show that an isolated cell can experience temporally complex alternans ranging from the resolution of AP alternans over a minute to complex amplitude varying, and even unstable alternans leading to loss of 1:1 capture. Further, the data demonstrate that alternans can be initiated in the absence of CaT induced by BAPTA. Lastly, consistent with previous reports that the I_CaL_ agonist and inotropic agent Bay k 8644 increases the magnitude of alternans,^30, 31^ we demonstrate that Bay k 8644 in the presence of BAPTA enhances the development of and destabilizes AP alternans.

Our data provides a revised interpretation on the discussion of whether CaT drives AP alternans and reveals a potentially new pro-arrhythmic scenario for the development of unstable (aka crescendo) AP alternans and activation block.

## METHODS

All procedures were approved by the Animal Care and Use Committee of the University of Utah and complied with the American Physiological Society’s *Guiding Principles in the Care and Use of Animals.* Expanded methods can be found in the **online supplement/appendix**.

### Myocyte isolation and electrophysiological methods

Adult New-Zealand White rabbit (1.5-1.7 Kg) ventricular myocytes were isolated by combining enzymatic tissue digestion and mechanical trituration (supplement for details).^32^ Transmembrane voltage (V_m_) was continuously recorded using a glass pipette (2-4 MΩ tip) and an Axoclamp 2B amplifier (Molecular Devices, Ca) in bridge mode (supplement for details).^33^ To elicit an AP, we injected a rectangular depolarizing current pulse (duration 3-4 ms) into the myocytes through the patch pipette (supplement for details). Using a previously described approach we loaded the fluorescent probe fluo-4 into the isolated myocytes to track the changes in intracellular Ca^2+^.^32, 34^ Fluo-4 fluorescence signals were recorded using an EMCCD camera (iXon 860, Andor Technology, Belfast, UK) configured to record images at a resolution of 64×64 pixels and 860 frames/s (supplement for details).^35^ Simultaneously recorded APs and CaTs were aligned for analysis (supplement for details) (**Supplement Fig. 1**). Where indicated, myocytes were loaded through the glass pipette with 10.0 mM or 20.0 mM BAPTA (1,2-Bis(2-aminophenoxy)ethane-N,N,N’,N’-tetraacetic acid; Tocris Bioscience, UK),^32^ a Ca^2+^ buffer used to effectively eliminate the CaT. Experiments were carried out at 36.5±1.0 °C using Rod-shaped myocytes with well-defined striations.

Once a myocyte was attached to the patch pipette, stimulus pulses were delivered at a pacing cycle length (BCL) of 2000 (or 1000) ms for 2-7 min to establish steady state conditions. Therein, the cells were subject to progressively shorter BCL with the objective of inducing AP alternans (**Supplement Fig. 2A**). The target BCLs were (in ms): 2000, 1000, 500, 300, 180. Given that each cell responded uniquely to changes in rate, a number of cells were subject to intermediate BCL during implementation of the pacing protocol. At each test BCL (including intermediate rates) the cells were constantly paced for 30 to 90 seconds before increasing the rate (**Supplement Fig. 2A**). Ultimately, the BCL was reduced to attain a final value of 180 ms unless the development of alternans occurred before.

### Experimental protocols

The pacing protocol (**Supplement Fig. 2A**) was applied to myocytes from four experimental groups: ***I*)** myocytes patched with normal pipette solution and superfused with normal Tyrode solution, *<u>Control group</u>* (n=22); ***II*)** cells patched with normal pipette solution and superfused with Tyrode’s solution containing 25 nM Bay K 8644, *<u>Bay K 8644 group</u>* (n=9); ***III*)** cells patched with pipette solution containing 10 mM BAPTA and superfused with normal Tyrode solution, *<u>10</u> <u>mM BAPTA group</u>* (n=10); ***IV*)** cells patched with pipette solution containing 20 mM BAPTA and superfused with normal Tyrode solution, *<u>20 mM BAPTA group</u>* (n=9). In the Bay K 8644 group, the pacing protocol was initiated once delivery of Bay K 8644 at baseline elicited a visible effect on the AP (i.e. AP prolongation).

To test the effect of Bay K 8644 on myocytes lacking CaT, myocytes loaded with 10 mM BAPTA bathed in control solution were exposed to 25nM Bay K 8644 either during constant pacing at 500-1000 ms BCL (n=8), or once 300-350 ms BCL was established via progressive BCL shortening (n=11). The BCL was adjusted to ensure 1:1 capture at the fastest possible rate as required by the drug effects.

#### Detection of APD alternans

Data reported were generated from the last 40 activations recorded at each test BCL (**Supplement Fig. 2A**). For each AP in this window we determined the APD at 90% repolarization (APD_90_)(supplement for details). Representative AP’s and APD_90_ sequences during 2:2 stimulus-responses are shown in **Supplement Figure 2B** (upper and lower, respectively). For each 40 activation-long analysis window we determined the maximum number of consecutive APD_90_ pairs locked in a 2:2 pattern. If 7 or more consecutive APD_90_ pairs exhibiting 2:2 fluctuations were identified, we designated the sequence as containing consolidated alternans and classified the myocytes as ‘alternans positive’, following the probabilistic reasoning introduced by Dilly et al. ^36^ The longest applied BCL satisfying the alternans positive criteria was defined as the ‘alternans threshold’. Myocytes devoid of APD_90_ sequences satisfying the ‘alternans positive’ criteria were classified as ‘alternans free’.

#### Data and statistical analysis

Data are presented as mean±standard deviation. For each test BCL, we computed the mean DI (Mean-DI) and mean APD_90_ (Mean-APD_90_) as the average DI and APD_90_ values of activations conforming the 40-activation-long analysis widow when consolidated alternans were absent, or across the duration of the consolidated alternans sequence when it was present. Myocytes where classified into either an ‘alternans positive” or ‘alternans free’ category, and the segregated data used for group-wise and/or category-wise statistical analysis by means of the Student’s t-test. We used the Chi-squared test to compare the group-wise fraction of alternans positive myocytes and the group-wise distribution of alternans types. Statistical comparisons yielding P values less than 0.05 were considered significant.

## RESULTS

### AP and CaT alternans

Action potential (AP) alternans were frequently induced in some but not all myocytes with rapid pacing. Representative AP and CaT traces in AP-alternans positive cells and AP-alternans negative cells can be seen in Figure 1. There were no significant differences between the fraction of AP alternans inducible cells between control (64%), cells loaded with the L-type calcium current (I_Ca,L_) agonist Bay K 8644 (25nM, 89%), or cells loaded with 10mM or 20mM Bapta (70% and 89% respectively) to abolish CaT.

**Figure 1.**
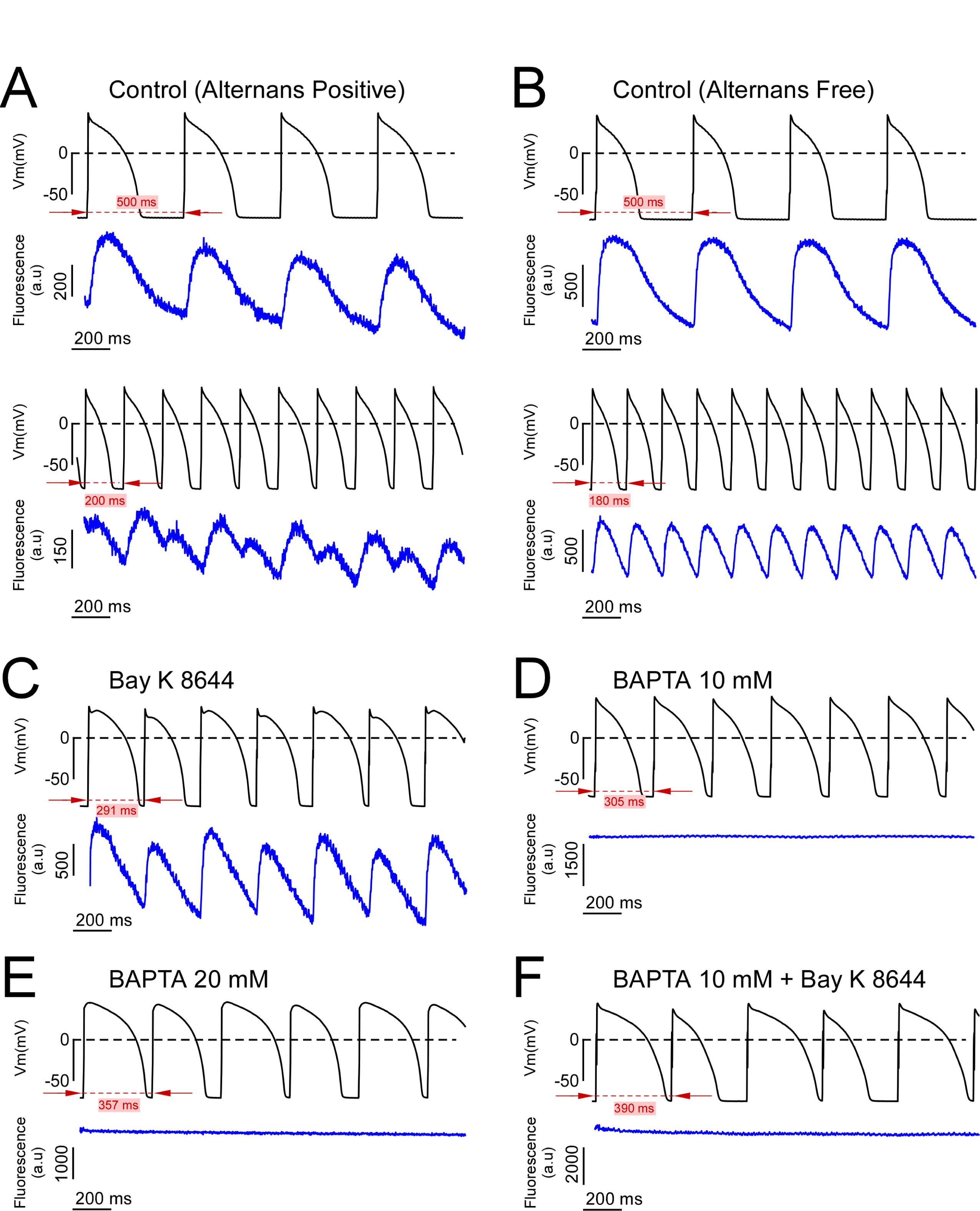
Dual AP-CaT recordings from alternans positive and alternans free cells. **A-B**. Simultaneous AP (upper trace, black) and CaT (lower trace, blue) recordings acquired during slow (upper pair of traces) and rapid (lower pair of traces) pacing from representative control alternans positive (**A**) and alternans negative (**B**) myocytes. **C-F**. Simultaneous AP (upper trace, black) and Fluo-4 fluorescence signals (lower trace, blue) recorded at (or below) the alternans threshold of alternans positive myocytes from the Bay K 8644 (**C**), 10 mM BAPTA (**D**), 20 mM BAPTA (**E**), and 10 mM BAPTA plus Bay K 8644 (**F**) groups.

Simultaneous AP and CaT recordings from a representative alternans positive myocyte illustrate how shortening the BCL from 500 ms to a value below the alternans threshold (BCL=220 ms) shifted the 1:1 captured beats to a dynamic 2:2 capture in both the AP and the CaT signals (Fig 1A). Due to the finite and relatively short optical CaT recordings relative to continuous voltage clamp recordings, we could not determine if either the AP or CaT alternans preceded one another, or if alternans occurred simultaneously in both signals. All dual AP and CaT recordings acquired during the development of AP alternans (12 dual recordings from 8 different cells) revealed an in-phase 2:2 capture in the AP and CaT signals, such that long APD was accompanied by large amplitude CaT, and conversely short APD with a small amplitude CaT (Fig 1A, lower two traces). Simultaneous AP and CaT recordings from a representative alternans free myocyte depict the absence of AP alternans in the 500-180 ms range of test BCLs (Fig 1B). Note that the simultaneously recorded CaT signals were also alternans free (Fig 1B, blue traces). In this study, CaT alternans were not observed in the absence of AP alternans. As expected, AP and CaT alternans were observed in the presence of the I_Ca,L_ agonist Bay K 8644 (Fig 1C). Interestingly, AP alternans were inducible in the absence of CaT secondary to perfusion of either 10 or 20mM BAPTA (Fig 1D-E). Importantly, agonizing I_Ca,L_ in the absence of CaT, further induced complex AP alternans (Fig 1F).

### Dynamic response of AP alternans

Analysis of the AP dynamic response to a pacing rhythm change at the scale of tens-to-hundreds of sequential activations revealed that the pattern of developing alternans was pleomorphic, in line with previous observations.^36^ We classified AP alternans into three types: overdamped, critically damped, and underdamped (Fig 2). Overdamped AP alternans were present at a rhythm change, and disappeared (aka resolved) after a variable amount of time. APD_90_ is plotted against time for a single pacing rate in Figure 2A-C for the purpose of visualizing the dynamic response of alternans over time. A representative trace of overdamped AP alternans is shown in Fig 2A.

**Figure 2.**
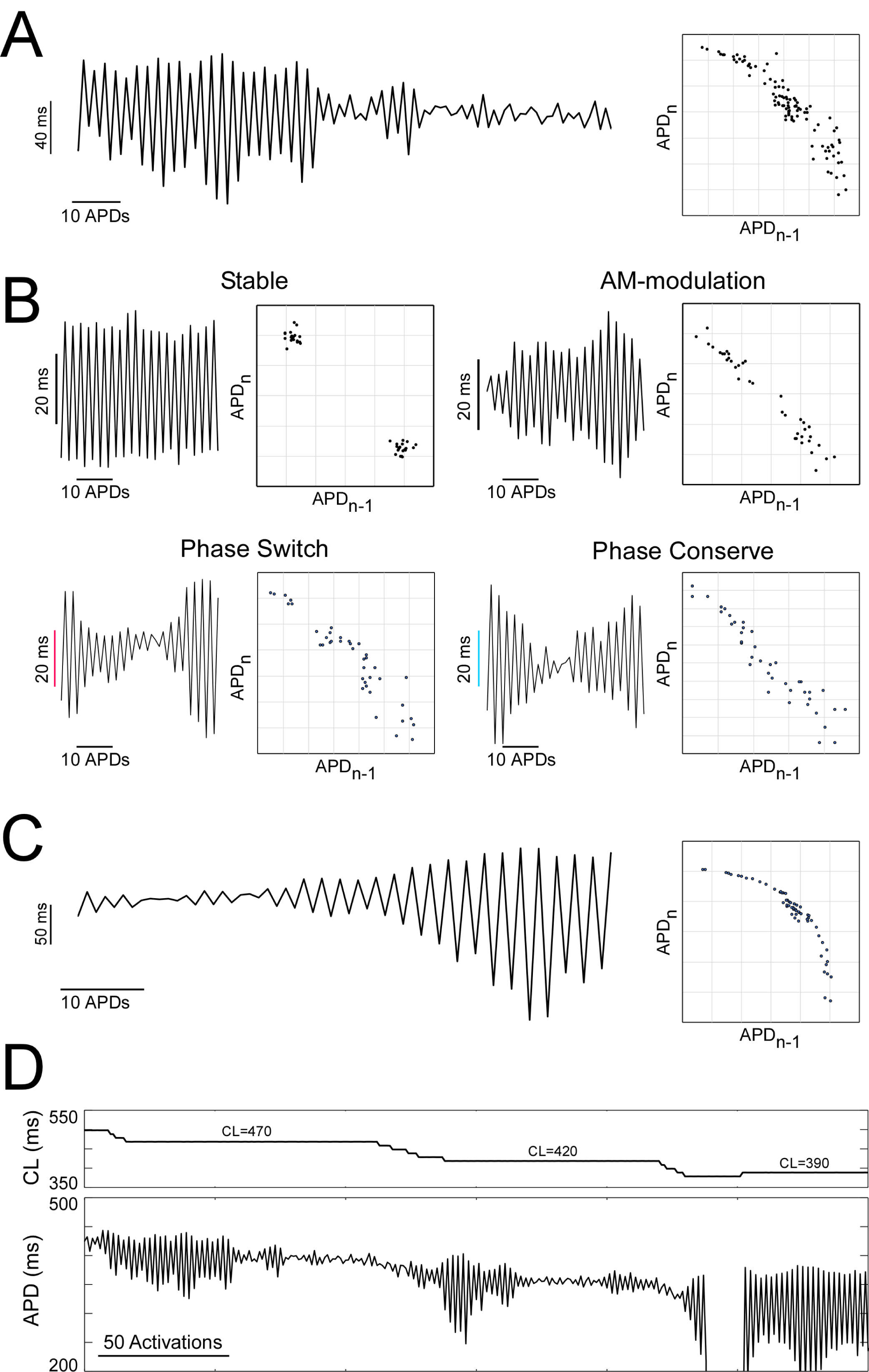
Types of dynamic response of AP alternans. **A-C**. APD_90_ values plotted against activation number for representative cases of overdamped (**A**), critically dampened (**B**), and underdamped (**C**) AP alternans. **Panel B** shows the four categories of critically damped alternans recorded: ‘stable’, ‘amplitude modulated’, ‘phase conserve, and ‘phase switch’. Each APD_90_ sequence in **panels A-C** has attached the corresponding Poincare plot constructed by recursively plotting a given APD_90_ (APD_n_) against the preceding APD_90_ (APD_n-1_). **D**. APD_90_ sequence (lower panel) exhibiting mixed AP alternans behavior in response to changes in the applied BCL (upper panel). APD_90_ values and BCL are plotted against the activation number.

Critically damped alternans exhibited at least 4 different behaviors: Resolution to stable alternans, alternans with lower frequency harmonics (amplitude modulated), alternans that resolved for some period of time into a brief alternans free sequence, and returned in-phase with the original sequence (AB - no alternans - AB), and finally alternans that resolved for some period of time and returned out of phase (AB - no alternans - BA). Representative data of the 4 critically damped AP alternans patterns (‘stable’, ‘amplitude modulated’, ‘phase conserve’, and ‘phase switch’) are shown in Fig 2B. An expanded analysis of examples exhibiting ‘phase conserve’ and ‘phase switch’ AP alternans sequences is shown in **Supplement Figure 3**.

Underdamped alternans, also known as crescendo alternans,^36^ were those that became progressively larger until 2:2 capture converted to 2:1 capture (Fig 2C).

Importantly, a single myocyte often exhibited a number of AP alternans behaviors during decremental and incremental pacing as can be seen Fig. 2D. To account for the highly variable nature of AP alternans, APs from the last 40 activations at a cycle length, with at least 7 consecutive 2:2 alternan pairs were quantified. This approach allowed for comparison of AP alternans thresholds (ALT-TH), APD_90_ and DI at the ALT-TH, and number of alternating pairs within the last 40 beats.

Analysis of summary (Table 1) reveals that AP alternans were more frequently critically damped in alternans positive cells of both the Control (77%) and the Bay K 8644 (43.75%) groups, with the proportion of underdamped alternans increasing from 9% to 31.25% in the Bay K 8644 group. Notably, the lower 10mM BAPTA concentration was associated to a reduced proportion of critically damped alternans (41.6%) occurring at the expense of an increase in overdamped alternans (41.6%)(Table 1), that resolved to non-alternating rhythms during rapid pacing episodes in 5/7 alternans positive myocytes. This finding is consistent with other reports that AP alternans are absent at steady-state pacing in the presence of BAPTA.^27^ Importantly, the AP alternans in 7/8 alternans positive cells treated with 20mM BAPTA were either critically damped, consolidated, and persistent (43.75%), or underdamped leading to crescendo alternans (43.75%)(Table 1).

**Table 1.** Group wise distribution of critically damped, overdamped, and undamped type alternans

|  | **Critically Damped (%)** | **Overdamped (%)** | **Underdamped - Crescendo (%)** | **Chi-square p value** |
| --- | --- | --- | --- | --- |
| **Control** | 77 | 14 | 9 |  |
| **Bay K 8644** | 43.75 | 25 | 31.25 | 0.046 |
| **10 mM BAPTA** | 41.6 | 41.6 | 16.6 | 0.069 |
| **20 mM BAPTA** | 43.75 | 12.5 | 43.75 | 0.012 |
| **BAPTA + Bay K 8644** | 7 | 14 | 79 | <0.0001 |

These data demonstrate that agonizing I_Ca,L_, or abolishing CaT can enhance the propensity for dynamic AP alternans, suggesting a negative feedback mechanism may exist between CaT and I_Ca,L_ to dampen AP alternans.

### AP alternans characteristics

Representative AP alternans and a quantification of APD_90_ over time for two cycle lengths are presented in Figure 3 for untreated control cells or cells treated with Bay K 8466 or the calcium chelator BAPTA.

**Figure 3.**
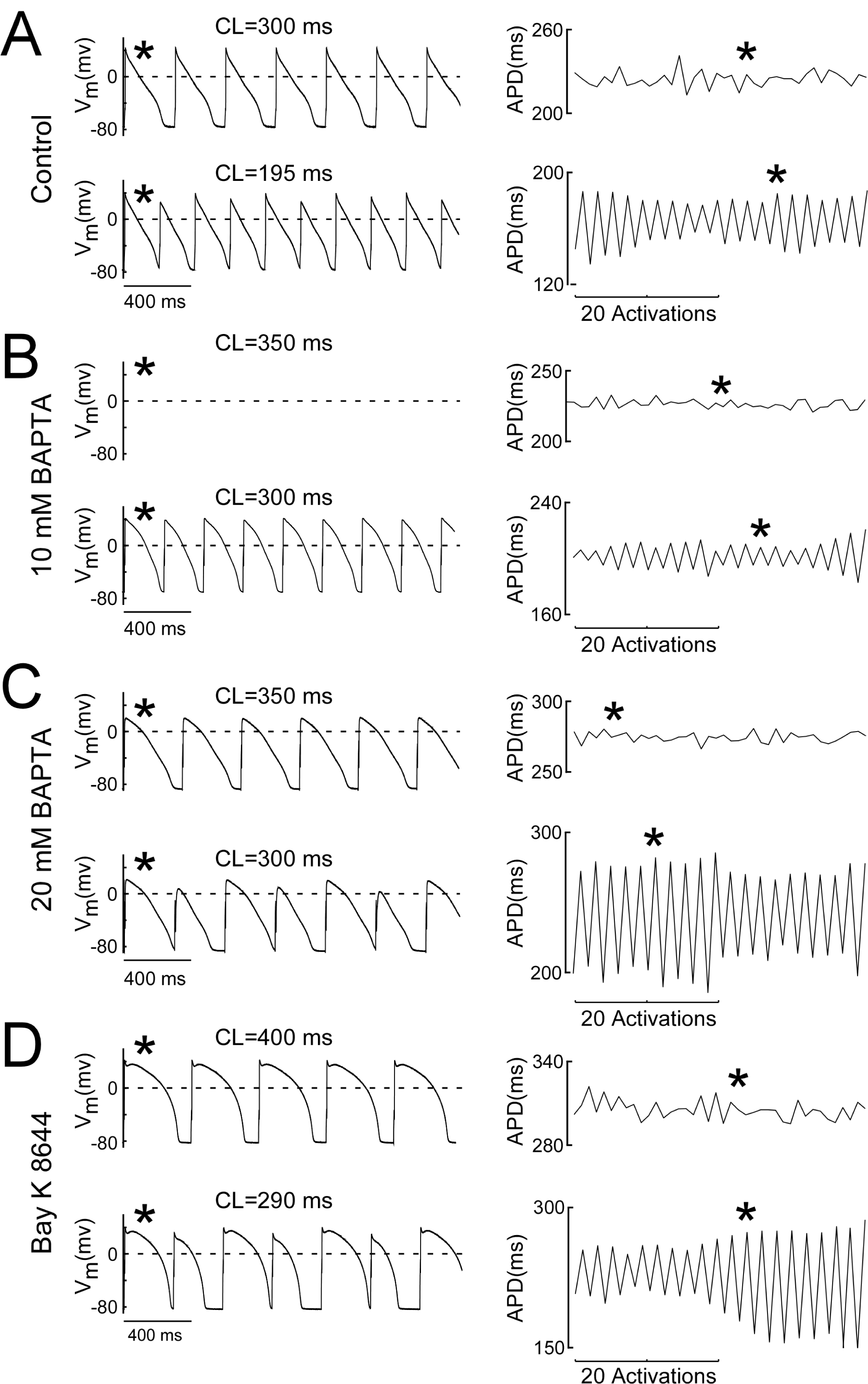
AP alternans prevail in Ca^2+^ buffered cells. **A-D**. Leftmost pairs of traces depict APs recorded during constant pacing before the onset of alternans (upper trace) and at the alternans threshold (lower trace depicting alternans) in control myocytes (**A**), myocytes loaded with 10 mM BAPTA (**B**), myocytes loaded with 20 mM BAPTA (**C**), and myocytes superfused with 25 uM Bay K 8644 (**D**). Rightmost pairs of traces in each panel depict the 40 activation-long APD_90_ sequences used to evaluate the presence of alternans at the corresponding test BCL. The asterisk indicates the correspondence between the AP and the APD_90_ measure. Labels above the AP recordings indicate the BCL.

Summary data in Table 2 demonstrates that Mean-APD_90_ at 500ms BCL (normocardia) and in the absence of AP alternans was 252±53 ms in controls. Whilst normocardic Mean-APD_90_ values in the Bay K 8466 group (296±65 ms) were not significantly different from the control group (Table 2), perfusion of the agonist to untreated myocytes during bradycardic pacing elicited APD_90_ prolongation from 270±107 ms to 323±116 ms (p<0.0001). Both 10 mM BAPTA and 20 mM BAPTA increased normocardic Mean-APD_90_ to 367±56 ms and 337±43 ms respectively (p<0.0005, Table 2).

**Table 2.** AP alternans characteristics

| Group | BCL = 500 ms |  | Group | BCL = ALT-TH |  |  |  |  |
| --- | --- | --- | --- | --- | --- | --- | --- | --- |
|  | DI (ms) | APD<sub>90</sub> (ms) |  | ALT-TH (ms) | DI (ms) | APD<sub>90</sub> (ms) | Number 2:2 AP's | ALT-MAG (ms) |
| **Control**<br>(n=22) | 248±51 | 252±53 | **Control**<br>(n=14) | 229±36 | 47±17 | 182±34 | 28±10 | 17±15 |
| **BAPTA10**<br>(n=10) | 130±55 ‡ | 367±56 ‡ | **BAPTA10**<br>(n=7) | 288±91 * | 49±13 | 240±79 * | 24±9 | 13±6 |
| **BAPTA20**<br>(n=9) | 160±43 ‡ | 337±43 ‡ | **BAPTA20**<br>(n=8) | 284±50 † | 53±11 | 232±44 † | 24±11 | 21±22 |
| **Bay K 8644**<br>(n=9) | 201±65 * | 296±65 | **Bay K 8644**<br>(n=8) | 262±72 | 56±14 | 206±60 | 27±11 | 30±38 |

\*p<0.05, Student's t-test, compared to control values of the same BCL category.
†p<0.01, Student's t-test, compared to control values of the same BCL category.
‡p<0.0005, Student's t-test, compared to control values of the same BCL category.
AP: Action Potential
ALT-TH: Alternans Threshold
ALT-MAG: Alternans Magnitude
APD<sub>90</sub>: Action Potential Duration at 90% repolarization
DI: Diastolic Interval
BCL: Pacing Cycle Length

In control, AP alternans positive cells, the mean BCL at which alternans occurred (the alternans threshold) was 229±36 ms (Table 2). Consistent with the lack of effect of 25 nM Bay K 8466 on normocardic Mean-APD_90_ values, the I_CaL_ agonist did not significantly increase mean alternans threshold (262±72 ms) relative to control (Table 2). In contrast, AP alternans were induced at slower rhythms in the absence of the CaT. Specifically, the AP alternans threshold in cells perfused with 10mM and 20mM BAPTA (288±91 ms and 284±50 ms respectively) were both significantly (p<0.05) greater than the threshold for control myocytes (Table 2).

Despite the complexity of alternans, we next quantified the number of consecutive APD_90_ pairs locked in a 2:2 pattern (Fig. 3A-D). Importantly, over a 40 beat window, we did not detect a significant difference in the number of consecutive 2:2 AP’s in control conditions or in cells treated with Bay K 8466, 10mM BAPTA, or 20mM BAPTA (Table 2), nor did we detect group-wise differences in the alternans magnitude measured at the ALT-TH (Table 2).

In order to compare Mean-APD_90_ between conditions associated with different alternans thresholds, the average Mean-APD_90_ at the alternans threshold, independent of BCL, can be seen in Table1. Interestingly, despite the Mean-APD_90_ prolongation elicited by Bay K 8466 during bradycardic pacing and consistent with the lack of effect during normocardia, the average Mean-APD_90_ at the alternans thresholds for individual myocytes was not significantly different under control conditions (182±34 ms) or in cells perfused with 25nm Bay K 8466 (206±60 ms)(Table 2). Similarities between these Mean-APD_90_ values were consistent with the similar alternans threshold values recorded for both groups (Table 2). In contrast, the average Mean-APD_90_ at the alternans thresholds for individual myocytes was significantly increased by 10nM BAPTA (240±79 ms) and 20nM BAPTA (232±44 ms), consistent with the higher alternans threshold observed in both CaT ablated groups (Table 2).

Interestingly, the average Mean-DI at the alternans thresholds was similar between interventions. Specifically, average Mean-DI at control was 47±17 ms and 56±14 ms with 25nM Bay K 8466, 49±13 ms with 10nM BAPTA, and 53±11 ms with 20nM BAPTA (Fig 3F), suggesting that a critical time-dependent event occurring during the DI may underlie alternans formation.

Together, the enhanced cellular susceptibility to alternans (higher alternans threshold) resulting from CaT removal indicates that CaT dampens alternans, likely via a negative feedback mechanism involving Ca^2+^ dependent I_Ca,L_ inactivation. Given that Bay K 8644 (previously reported by Valderrabano et al.^31^ to promote alternans at 100 nM) did not significantly promote the formation of alternans when delivered at 25 nM, we reasoned that ‘protection’ against alternans development may be conferred by the presence of CaT.

### Combined CaT elimination and agonizing I_Ca,L_ promotes crescendo alternans

As a final exploration of the feedback loop between AP, CaT, and I_Ca,L_, cells were first perfused with the lowest BAPTA concentration (10mM) and then simultaneously perfused with 10mM BAPTA and 25nM Bay K 8466. We hypothesized that the agonizing I_Ca,L_ in the absence of CaT would further increase cellular susceptibility to AP alternans as well as the complexity of those alternans. The results were unexpected, yet still consistent with the hypothesis. Specifically, perfusion of Bay K 8644 upon myocytes lacking CaT prompted prominent changes in the dynamic response to pacing (Fig. 4). During bradycardic constant pacing (1000ms BCL), AP alternans were not observed with 10mM BAPTA as discussed previously (Fig. 4A-B). However, the addition of Bay K 8644 to CaT ablated myocyte (Fig. 4A-B, dotted line and bar initiating at stimulus #267) elicited a fast and prominent increase of APD_90_ (from ∼500 ms at baseline to ∼850 ms) followed by the formation of AP alternans (Fig. 4B, stimulus# 284 to 302). Of note, the magnitude of the alternans progressively increased and became unstable (Fig 4B, see trace contained inside the leftmost dotted-line box and the corresponding zoomed-in APD_90_ sequence displayed in the adjacent inset). The unrestricted increases in the magnitude of alternans during constant pacing ultimately lead to activation block (i.e. 2:1 capture) on stimulus# 303 (Fig 4B, leftmost blue asterisk). Attempts to abate the development of crescendo alternans by continuously readjusting the BCL to larger values (Fig 4A, solid line) proved unsuccessful as recurrent episodes of crescendo alternans leading to block occurred in succession (Fig. 4B, APD_90_ sequences contained in the 2^nd^, 3^rd^, and 4^th^-from left dotted-line boxes and the corresponding zoomed-in APD_90_ sequences displayed in the adjacent insets). Each asterisk upon the APD_90_ trace marks the point at which the diverging 2:2 activation converted to a 2:1 type response. The final events leading to the activation block highlighted by the red asterisk in the 2^nd^ from left dotted-line box, are labeled ‘a’ to ‘j’ in the corresponding zoomed-in APD_90_ sequence, and can be traced to the corresponding V_m_ changes depicted in Fig 4C. Note here that the progressive increase in the magnitude of AP alternans leads to the final abortive response to pacing which effectively initiates the 2:1 activation sequence (Fig 4C, red asterisk).

**Figure 4.**
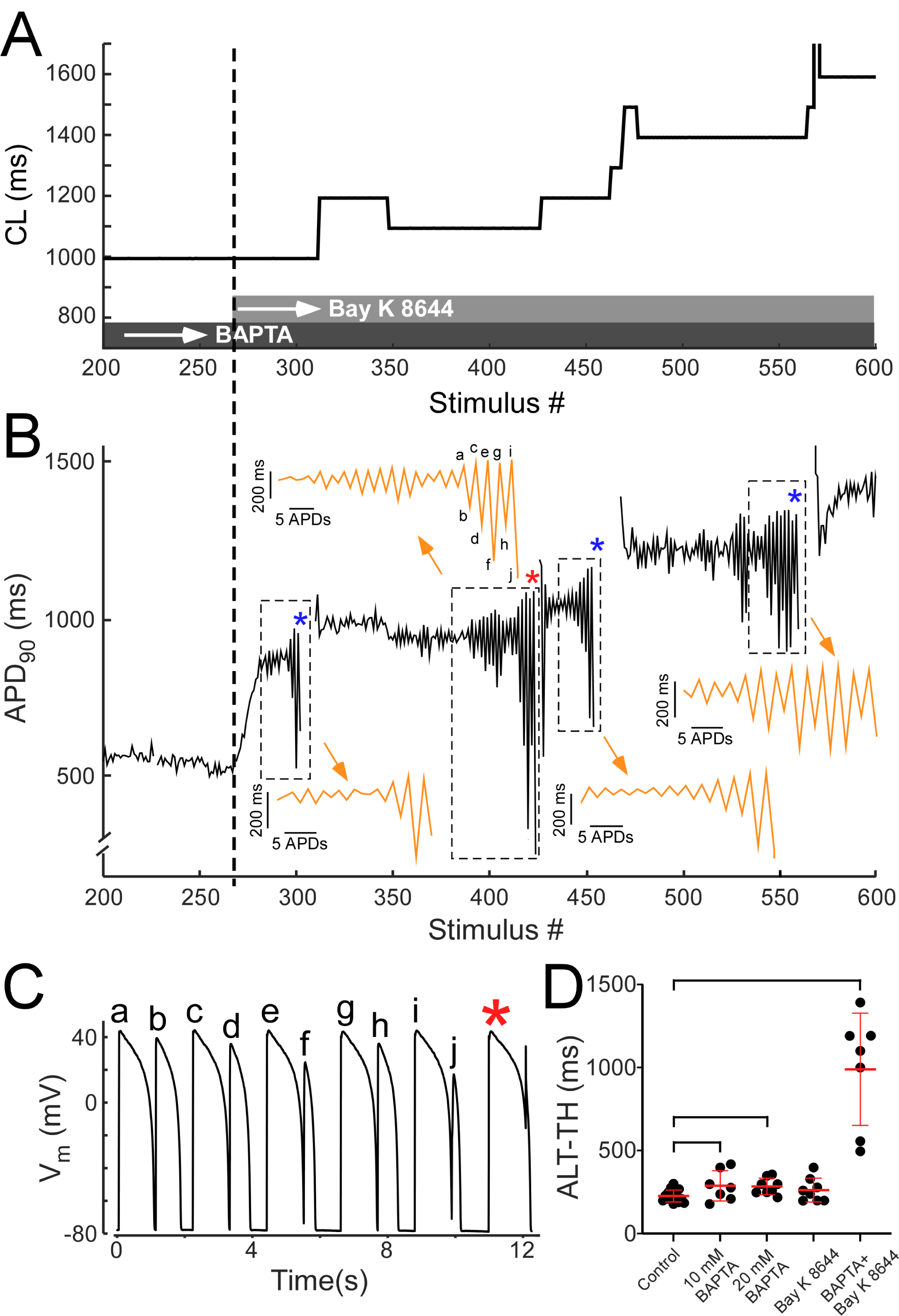
CaT ablation plus Bay K 8644 elicits unrestrained crescendo AP alternans. **A-B**. Time-course of applied BCL values (**A**) and APD_90_ changes (**B**) recorded from a representative CaT ablated myocyte subject to constant pacing (stimulus #200 through #600) during baseline and after perfusion with 25 nM bay K 8644. Dark grey and light grey bands overlying the abscissa in panel A identify the intervals in which 10 mM BAPTA and 25 uM Bay K 8644 were present. The dotted line spanning panels A-to-B traces the stimulus number at time of initiation of Bay K 8644 perfusion to the respective BCL and APD_90_ changes depicted in each panel. Dotted-line boxes in panel B identify four episodes of crescendo alternans. Adjacent to each box the orange traces depict the zoomed-in APD_90_ sequence corresponding to the crescendo alternans episode contained in the box. Asterisks identify the point in the APD_90_ trace at which the diverging 2:2 activation converted to a 2:1 type response. Sections containing APD_90_ values associated to 2:1 responses were deleted for clarity. Labels a-to-j identify the final APD_90_ values leading to 2:1 activation identified by the red asterisk. **C**. V_m_ changes exhibiting the transition from a 2:2 to a 2:1 activation from example identified by red asterisk in **panel B**. Labels a-to-c trace the correspondence between each AP in **panel C** and the APD_90_ value in **panel B**. **D**. Group-wise comparison of the alternans threshold. Black circles represent the individual alternans threshold values and red lines combined with error bars represent the alternans threshold mean and standard deviation values for control, Bay K 8644, 10 mM BAPTA, 20 mM BAPTA, and 10 mM BAPTA + Bay K 8644 groups. Brackets identify comparisons between average values exhibiting significant differences (p<0.05, Student t-test).

In all, similar type of responses were obtained from six additional myocytes subject to combined 10 mM BAPTA and Bay K 8644. In these myocytes, the proportion of underdamped crescendo alternans increased prominently from 9% in control to 79% (Table 1). Further, the alternans threshold in the CaT ablated myocytes subject to Bay K 8644 was markedly increased to 990±338 ms (p<0.0002 compared to control)(Fig 4D). Similar experiments carried out at physiological rates (BCL=300 ms) resulted in qualitatively similar development of crescendo alternans and block after Bay K 8644 delivery (**Supplement Fig. 4**). The longest BCL sustaining AP alternans in this group of cells was also significantly larger than controls (417±97 ms p<0.005 compared to control). The data therefore demonstrate that in the absence of CaT, agonizing I_Ca,L_ is proarrhythmic at a cellular level.

## DISCUSSION

Our study is the first report demonstrating an AP alternans promoting effect of the Ca^2+^ chelator BAPTA, used to eliminate the CaT. Further, we show that CaT ablation in combination with the I_CaL_ agonist and inotropic agent Bay K 8644 can promote the formation of crescendo alternans and subsequent activation block, an arrhythmia triggering event.

### The data are clinically relevant because they provide mechanistic insight into the origin of AP alternans, the cellular driver of TWA, a known harbinger of ventricular arrhythmia

Our results are in contrast with those of Walker et al.^28^ and of Goldhaber et al.^27^ showing that CaT removal (with BAPTA) protected against the development of alternans.^11, 23–25, 27, 28^ Discrepancies with Walker et al.’s work may originate in their experimental temperature settings (26 °C), a condition well known to modify the development alternans, as well as in their use of BAPTA at much lower concentrations which reportedly failed to change APD.^28^ Experimental conditions between the study of Goldhaber et al. and ours were comparable, yet in contrast to our data, the authors reported a complete elimination of AP alternans with 0.1-10 mM BAPTA (also with BAPTA-AM, and with Thapsigargin plus BAPTA). Whilst the results of both studies are nominally opposite (protective vs detrimental), the authors also reported a BAPTA-induced APD shortening occurring at rapid pacing rates which may underlie the contrasting outcomes between studies. Of note, whilst our observation that 10 mM BAPTA promoted the formation of ‘overdamped’ type alternans may be explained by a buffer-induced APD shortening similar to that reported by Goldhaber et al., our observation that 20 mM BAPTA promoted ‘underdamped’ crescendo alternans is incompatible with a rate dependent APD shortening effect of BAPTA.

Towards determining how CaT elimination modulates the formation of alternans, others used either caffeine,^11, 23, 24^ ryanodine,^11, 23, 25^ thapsigargin,^26^ ryanodine in combination with thapsigargin,^27^ as a strategy to remove CaT. All listed approaches prevent sarcoplasmic reticulum Ca^2+^ accumulation, which should increase cellular free Ca^2+^ in contrast to the buffering effect of BAPTA on cytosolic Ca^2+^ homeostasis. The contrasting protective vs detrimental outcomes of ablating CaT when comparing data-sets may result from these distinct experimental approaches. It is plausible that differing free Ca^2+^ levels between approaches may result in distinct effects on the Ca^2+^-dependent inactivation of I_CaL_, with approaches increasing the cytosolic Ca^2+^ levels promoting I_CaL_ inactivation (protective) versus those approaches reducing cytosolic Ca^2+^ levels preventing I_CaL_ inactivation (detrimental). In line with this notion, Sham et al. showed that 10 mM BAPTA reduced Ca^2+^-dependent inactivation of I_CaL_ in pulmonary vein myocytes.^37^ Additional studies addressing how I_CaL_ gating is modulated by the interaction of BAPTA, Bay k 8644, and activation rate will be required to further understand the role of the Ca^2+^-dependent inactivation of I_CaL_ in promoting alternans.

Aside of the origin of discrepancies amongst studies, our data prove that Ca^2+^ alternans are not required for the development of rapid-pacing induced AP alternans at physiological temperatures. Importantly, in the CaT ablated myocyte the alternans threshold is increased (i.e. slower rate required). This principle finding and must be viewed in tandem with earlier studies demonstrating the (close to) reverse scenario, that is the induction of Ca^2+^ alternans driven by an alternans-free voltage signal (AP-clamp).^21^ Taken together, the data indicate that both signals are capable of generating alternans inherently (i.e. in isolation). Thus, a principle question remains: which of both signals drives alternans in a first instance (i.e. is the precursor) when both signals are physiologically bi-directionally coupled. Indeed, there may not be a unique answer. Towards resolving this, comparing the alternans threshold of each signal in isolation would be useful, as it would reflect the inherent predisposition of each signal to generate alternans. Our data place the CaT ablated AP alternans threshold prominently higher (∼280 ms) than that of the bidirectionally coupled AP and CaT signals which is ∼220 ms, in accordance with previous studies.^38^ The equivalent measure for the CaT, the inherent Ca^2+^ alternans threshold, has not been, to the best of our knowledge measured in rabbit ventricular myocytes. The study by Chudin et al.^21^ provided a single experimental measure of AP-clamped Ca^2+^ alternans developing at 180 ms in rabbit myocytes and placed the Ca^2+^ alternans threshold value at 250 ms using simulations, but did not clarify if this was achieved under AP-clamp conditions or not. Others showed that in dog alternans free AP-clamped myocytes the threshold for Ca^2+^ alternans was ∼258 ms (of note, these measurements were performed at 32 °C).^39^ Side-to-side comparison of the available information on the inherent thresholds for the AP and the CaT signals suggests that AP signals are most inherently prone to develop alternans in response to rapid pacing (i.e. higher threshold). Given the above, our data is consistent with a model in which rapid-pacing induced AP alternans are governed by a voltage driven mechanism which is dampened by the presence of CaT via bi-directional coupling of AP and CaT signals. Given the prominent BAPTA-induced APD prolongation observed both at baseline and at the AP alternans threshold, it is likely that the bi-directional coupling mechanism involved in dampening AP alternans is a CaT mediated-inactivation of I_CaL_. Under this scenario, we envision the determinants of intracellular Ca^2+^ release restitution,^40^ as being key in modulating AP alternans through dynamic dampening of the AP oscillations. Experiments designed to measure the inherent capability of CaT to develop alternans at physiological temperatures as well as tandem measurements during transition to alternans will be required to further clarify the mechanism driving AP alternans. Towards this, simultaneous recordings of AP and CaT during rapid pacing induced alternans in intact rabbit hearts showed a temporal hierarchy in the development of alternans with CaT amplitude alternans leading followed by APD alternans.^41^ Others, however, reported the simultaneous initiation of both AP and CaT alternans in dog hearts.^39^ Given the limitations of optical mapping (limited spatial resolution plus imaging limited to the epicardial layers) experiments in isolated myocytes may be useful towards determining if any of the signals is the alternans precursor.

Our experiments also revealed the formation of crescendo alternans when we combined CaT ablation with low concentrations of Bay K 8644. It is noteworthy that the delivery of Bay K 8644 to myocytes with conserved CaT had a modest effect on the development of alternans, which is consistent with previous reports carried out using similar concentration of the drug.^30^ Remarkably, identical delivery of Bay K 8644 to CaT ablated myocytes had a starkly distinct (and arrhythmogenic) response consisting of the development of crescendo alternans followed by activation block. This occurred recursively at progressively slower pacing rates precluding implementation of 1:1 activation in CaT ablated myocytes, at rates much slower even than the those attained at baseline in CaT conserved myocytes. Again, the data support the notion that CaT dampens the development of AP alternans and precludes the initiation of activation block of what otherwise would be unconstrained oscillations of the transmembrane voltage, likely driven by fluctuations in the activation and deactivation of I_CaL_.

The notion that CaT provides an alternans protective effect via a bi-directional coupling mediated dampening mechanism has practical implications related to the delivery of therapy since various cardiac conditions are associated to reduced CaT. A case in point is heart failure, which has been associated to reductions of the CaT of up to 50%,^42^ or even beyond in terminally failing human hearts.^43^ In this scenario, it is plausible that reduced CaT may less efficiently ‘dampen’ (voltage-driven) AP alternans thus effectively promoting their development in the diseased heart (according to our data, this could be particularly damaging if therapy involved activating I_CaL_). However, if such a partial reduction of the CaT has a pro-AP alternans consequence remains to be investigated. Of note however, heart failure is indeed associated to an increased propensity to develop AP alternans, although the authors of the study attributed this to the increased propensity for inherent Ca^2+^ alternans to develop.^39^ Further studies will be required to determine how heart failure mediated CaT reductions alter the inherent ability of voltage driven AP alternans to develop.

Principle conclusions may be drawn from our study: 1) We demonstrate unequivocally that Ca^2+^ alternans are not required for the development of AP alternans; 2) Our data suggests that via bi-directional coupling, CaT (an possibly CaT alternans) provides a negative feedback signal that dampens the formation of rapid pacing induced AP alternans effectively reducing the threshold for their development in healthy hearts; 3) Therapies aiming to reduce CaT or its dynamic response, or alternatively inotropic agents targeting ICaL should consider the pro-alternans consequences of these interventions.

## Non-Standard Abbreviations and Acronyms

TWA: T-Wave Alternans
AP: Action Potential
CaT: Calcium Transient
ALT-TH: Alternans Threshold
ALT-MAG: Alternans Magnitude
APD: Action Potential Duration
APD_90_: Action Potential Duration at 90% repolarization
DI: Diastolic Interval
V_m_: Transmembrane Potential
BCL: Pacing Cycle Length
I_CaL_: L-type calcium current

## Acknowledgements

We would like to thank Drs. Natalia Torres, Kenneth W. Spitzer, and Alexey V. Zaitsev for their excellent technical and scientific advice.

## Supplemental Material

Supplemental Methods

Figures S1-S4

---

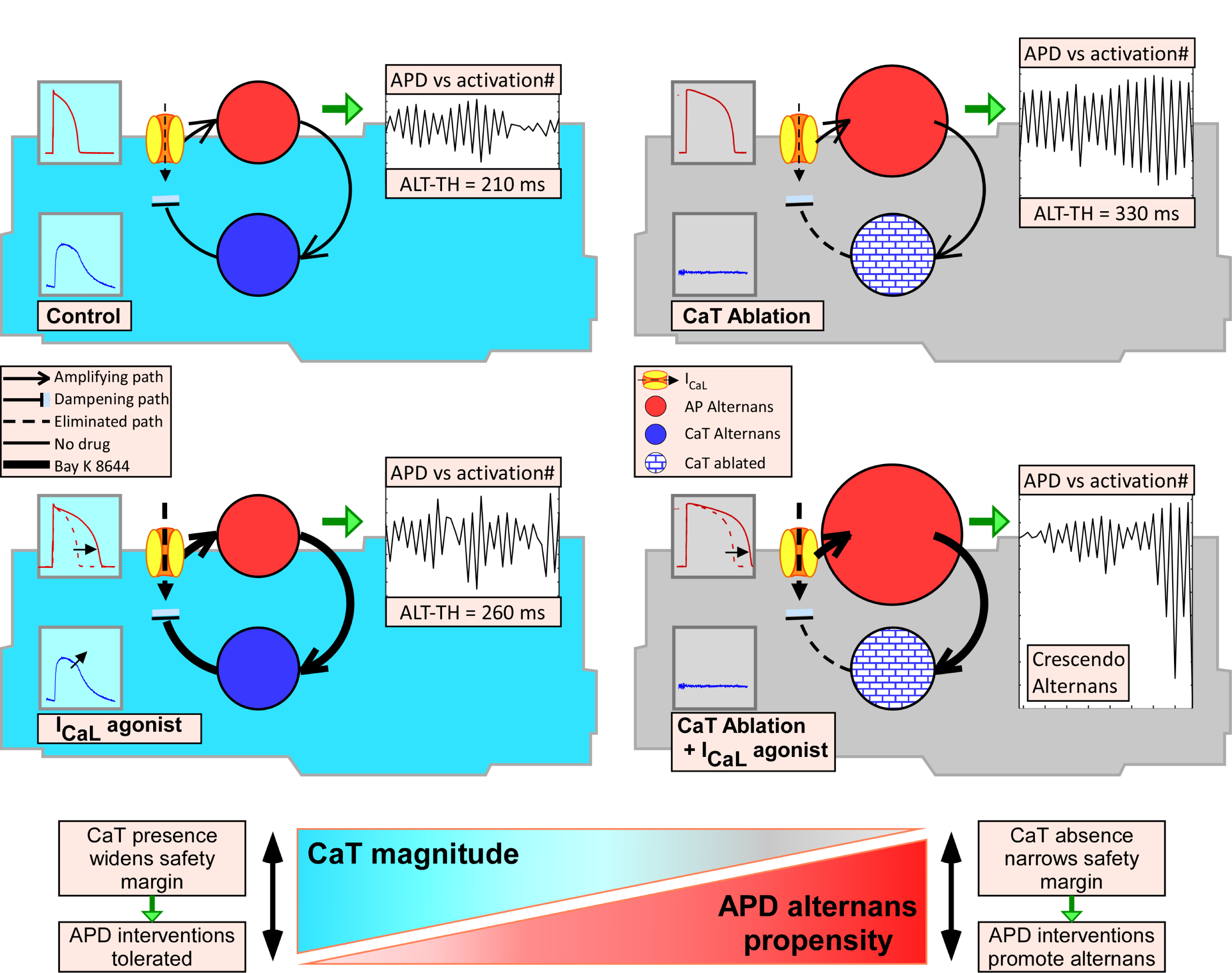

## REFERENCES

1. Cutler MJ, Rosenbaum DS. Explaining the clinical manifestations of T wave alternans in patients at risk for sudden cardiac death. Heart Rhythm. 2009;6:S22–28. doi: 10.1016/j.hrthm.2008.10.007

2. Verrier RL, Klingenheben T, Malik M, El-Sherif N, Exner DV, Hohnloser SH, Ikeda T, Martinez JP, Narayan SM, Nieminen T, et al. Microvolt T-wave alternans physiological basis, methods of measurement, and clinical utility--consensus guideline by International Society for Holter and Noninvasive Electrocardiology. J Am Coll Cardiol. 2011;58:1309–1324. doi: 10.1016/j.jacc.2011.06.029

3. Nearing BD, Verrier RL. Progressive increases in complexity of T-wave oscillations herald ischemia-induced ventricular fibrillation. Circ Res. 2002;91:727–732. doi: 10.1161/01.res.0000038887.17976.33

4. Shusterman V, Goldberg A, London B. Upsurge in T-wave alternans and nonalternating repolarization instability precedes spontaneous initiation of ventricular tachyarrhythmias in humans. Circulation. 2006;113:2880–2887. doi: 10.1161/CIRCULATIONAHA.105.607895

5. Narayan SM, Bode F, Karasik PL, Franz MR. Alternans of atrial action potentials during atrial flutter as a precursor to atrial fibrillation. Circulation. 2002;106:1968–1973. doi: 10.1161/01.cir.0000037062.35762.b4

6. Karma A. Electrical alternans and spiral wave breakup in cardiac tissue. Chaos. 1994;4:461–472. doi: 10.1063/1.166024

7. Garfinkel A, Chen PS, Walter DO, Karagueuzian HS, Kogan B, Evans SJ, Karpoukhin M, Hwang C, Uchida T, Gotoh M, et al. Quasiperiodicity and chaos in cardiac fibrillation. J Clin Invest. 1997;99:305–314. doi: 10.1172/JCI119159

8. Koller ML, Riccio ML, Gilmour RF, Jr. Dynamic restitution of action potential duration during electrical alternans and ventricular fibrillation. Am J Physiol. 1998;275:H1635–1642. doi: 10.1152/ajpheart.1998.275.5.H1635

9. Clusin WT. Mechanisms of calcium transient and action potential alternans in cardiac cells and tissues. Am J Physiol Heart Circ Physiol. 2008;294:H1–H10. doi: 10.1152/ajpheart.00802.2007

10. Lab MJ, Lee JA. Changes in intracellular calcium during mechanical alternans in isolated ferret ventricular muscle. Circ Res. 1990;66:585–595. doi: 10.1161/01.res.66.3.585

11. Kihara Y, Morgan JP. Abnormal Cai2+ handling is the primary cause of mechanical alternans: study in ferret ventricular muscles. Am J Physiol. 1991;261:H1746–1755. doi: 10.1152/ajpheart.1991.261.6.H1746

12. Euler DE. Cardiac alternans: mechanisms and pathophysiological significance. Cardiovasc Res. 1999;42:583–590. doi: 10.1016/s0008-6363(99)00011-5

13. Pastore JM, Girouard SD, Laurita KR, Akar FG, Rosenbaum DS. Mechanism linking T-wave alternans to the genesis of cardiac fibrillation. Circulation. 1999;99:1385–1394. doi: 10.1161/01.cir.99.10.1385

14. Pruvot EJ, Katra RP, Rosenbaum DS, Laurita KR. Role of calcium cycling versus restitution in the mechanism of repolarization alternans. Circ Res. 2004;94:1083–1090. doi: 10.1161/01.RES.0000125629.72053.95

15. Lee HC, Mohabir R, Smith N, Franz MR, Clusin WT. Effect of ischemia on calcium-dependent fluorescence transients in rabbit hearts containing indo 1. Correlation with monophasic action potentials and contraction. Circulation. 1988;78:1047–1059. doi: 10.1161/01.cir.78.4.1047

16. Qian YW, Clusin WT, Lin SF, Han J, Sung RJ. Spatial heterogeneity of calcium transient alternans during the early phase of myocardial ischemia in the blood-perfused rabbit heart. Circulation. 2001;104:2082–2087. doi: 10.1161/hc4201.097136

17. Weiss JN, Karma A, Shiferaw Y, Chen PS, Garfinkel A, Qu Z. From pulsus to pulseless: the saga of cardiac alternans. Circ Res. 2006;98:1244–1253. doi: 10.1161/01.RES.0000224540.97431.f0

18. Laurita KR, Rosenbaum DS. Cellular mechanisms of arrhythmogenic cardiac alternans. Prog Biophys Mol Biol. 2008;97:332–347. doi: 10.1016/j.pbiomolbio.2008.02.014

19. Qu Z, Xie Y, Garfinkel A, Weiss JN. T-wave alternans and arrhythmogenesis in cardiac diseases. Front Physiol. 2010;1:154. doi: 10.3389/fphys.2010.00154

20. Nolasco JB, Dahlen RW. A graphic method for the study of alternation in cardiac action potentials. J Appl Physiol. 1968;25:191–196. doi: 10.1152/jappl.1968.25.2.191

21. Chudin E, Goldhaber J, Garfinkel A, Weiss J, Kogan B. Intracellular Ca(2+) dynamics and the stability of ventricular tachycardia. Biophys J. 1999;77:2930–2941. doi: 10.1016/S0006-3495(99)77126-2

22. Kulkarni K, Merchant FM, Kassab MB, Sana F, Moazzami K, Sayadi O, Singh JP, Heist EK, Armoundas AA. Cardiac Alternans: Mechanisms and Clinical Utility in Arrhythmia Prevention. J Am Heart Assoc. 2019;8:e013750. doi: 10.1161/JAHA.119.013750

23. Saitoh H, Bailey JC, Surawicz B. Action potential duration alternans in dog Purkinje and ventricular muscle fibers. Further evidence in support of two different mechanisms. Circulation. 1989;80:1421–1431. doi: 10.1161/01.cir.80.5.1421

24. Hirayama Y, Saitoh H, Atarashi H, Hayakawa H. Electrical and mechanical alternans in canine myocardium in vivo. Dependence on intracellular calcium cycling. Circulation. 1993;88:2894–2902. doi: 10.1161/01.cir.88.6.2894

25. Kanaporis G, Blatter LA. The mechanisms of calcium cycling and action potential dynamics in cardiac alternans. Circ Res. 2015;116:846–856. doi: 10.1161/CIRCRESAHA.116.305404

26. Pearman CM, Madders GWP, Radcliffe EJ, Kirkwood GJ, Lawless M, Watkins A, Smith CER, Trafford AW, Eisner DA, Dibb KM. Increased Vulnerability to Atrial Fibrillation Is Associated With Increased Susceptibility to Alternans in Old Sheep. J Am Heart Assoc. 2018;7:e009972. doi: 10.1161/JAHA.118.009972

27. Goldhaber JI, Xie LH, Duong T, Motter C, Khuu K, Weiss JN. Action potential duration restitution and alternans in rabbit ventricular myocytes: the key role of intracellular calcium cycling. Circ Res. 2005;96:459–466. doi: 10.1161/01.RES.0000156891.66893.83

28. Walker ML, Wan X, Kirsch GE, Rosenbaum DS. Hysteresis effect implicates calcium cycling as a mechanism of repolarization alternans. Circulation. 2003;108:2704–2709. doi: 10.1161/01.CIR.0000093276.10885.5B

29. Shiferaw Y, Watanabe MA, Garfinkel A, Weiss JN, Karma A. Model of intracellular calcium cycling in ventricular myocytes. Biophys J. 2003;85:3666–3686. doi: 10.1016/S0006-3495(03)74784-5

30. Saitoh H, Bailey JC, Surawicz B. Alternans of action potential duration after abrupt shortening of cycle length: differences between dog Purkinje and ventricular muscle fibers. Circ Res. 1988;62:1027–1040. doi: 10.1161/01.res.62.5.1027

31. de Diego C, Pai RK, Dave AS, Lynch A, Thu M, Chen F, Xie LH, Weiss JN, Valderrabano M. Spatially discordant alternans in cardiomyocyte monolayers. Am J Physiol Heart Circ Physiol. 2008;294:H1417–1425. doi: 10.1152/ajpheart.01233.2007

32. Saegusa N, Garg V, Spitzer KW. Modulation of ventricular transient outward K(+) current by acidosis and its effects on excitation-contraction coupling. Am J Physiol Heart Circ Physiol. 2013;304:H1680–1696. doi: 10.1152/ajpheart.00070.2013

33. Zaniboni M, Pollard AE, Yang L, Spitzer KW. Beat-to-beat repolarization variability in ventricular myocytes and its suppression by electrical coupling. Am J Physiol Heart Circ Physiol. 2000;278:H677–687. doi: 10.1152/ajpheart.2000.278.3.H677

34. Saegusa N, Moorhouse E, Vaughan-Jones RD, Spitzer KW. Influence of pH on Ca(2)(+) current and its control of electrical and Ca(2)(+) signaling in ventricular myocytes. J Gen Physiol. 2011;138:537–559. doi: 10.1085/jgp.201110658

35. Warren M, Spitzer KW, Steadman BW, Rees TD, Venable P, Taylor T, Shibayama J, Yan P, Wuskell JP, Loew LM, et al. High-precision recording of the action potential in isolated cardiomyocytes using the near-infrared fluorescent dye di-4-ANBDQBS. Am J Physiol Heart Circ Physiol. 2010;299:H1271–1281. doi: 10.1152/ajpheart.00248.2010

36. Dilly SG, Lab MJ. Electrophysiological alternans and restitution during acute regional ischaemia in myocardium of anaesthetized pig. J Physiol. 1988;402:315–333. doi: 10.1113/jphysiol.1988.sp017206

37. Sham JS. Ca2+ release-induced inactivation of Ca2+ current in rat ventricular myocytes: evidence for local Ca2+ signalling. J Physiol. 1997;500 (Pt 2):285–295. doi: 10.1113/jphysiol.1997.sp022020

38. Mahajan A, Shiferaw Y, Sato D, Baher A, Olcese R, Xie LH, Yang MJ, Chen PS, Restrepo JG, Karma A, et al. A rabbit ventricular action potential model replicating cardiac dynamics at rapid heart rates. Biophys J. 2008;94:392–410. doi: 10.1529/biophysj.106.98160

39. Wilson LD, Jeyaraj D, Wan X, Hoeker GS, Said TH, Gittinger M, Laurita KR, Rosenbaum DS. Heart failure enhances susceptibility to arrhythmogenic cardiac alternans. Heart Rhythm. 2009;6:251–259. doi: 10.1016/j.hrthm.2008.11.008

40. Cely-Ortiz A, Felice JI, Diaz-Zegarra LA, Valverde CA, Federico M, Palomeque J, Wehrens XHT, Kranias EG, Aiello EA, Lascano EC, et al. Determinants of Ca2+ release restitution: Insights from genetically altered animals and mathematical modeling. J Gen Physiol. 2020;152. doi: 10.1085/jgp.201912512

41. Visweswaran R, McIntyre SD, Ramkrishnan K, Zhao X, Tolkacheva EG. Spatiotemporal evolution and prediction of [Ca(2+)]i and APD alternans in isolated rabbit hearts. J Cardiovasc Electrophysiol. 2013;24:1287–1295. doi: 10.1111/jce.12200

42. Piacentino V, 3rd, Weber CR, Chen X, Weisser-Thomas J, Margulies KB, Bers DM, Houser SR. Cellular basis of abnormal calcium transients of failing human ventricular myocytes. Circ Res. 2003;92:651–658. doi: 10.1161/01.RES.0000062469.83985.9B

43. Beuckelmann DJ, Erdmann E. Ca(2+)-currents and intracellular [Ca2+]i-transients in single ventricular myocytes isolated from terminally failing human myocardium. Basic Res Cardiol. 1992;87 Suppl 1:235–243. doi: 10.1007/978-3-642-72474-9_19

